# A triple-negative matrix-producing breast carcinoma is arrested by tumor-targeting *Salmonella typhimurium* A1-R in a PDOX model

**DOI:** 10.1101/2021.01.14.426747

**Authors:** Kazuyuki Hamada, Jun Yamamoto, Chihiro Hozumi, Ming Zhao, Takuya Murata, Norihiko Sugisawa, Yu Sun, Yusuke Aoki, Hiroto Nishino, Michael Bouvet, Takuya Tsunoda, Robert M. Hoffman

## Abstract

**Background/Aim:** Matrix-producing breast carcinoma (MPBC), is a rare, recalcitrant and highly aggressive. The present study aimed to determine the efficacy of tumor-targeting *Salmonella typhimurium* (*S. typhimurium*) A1-R on a triple-negative MPBC in a patient-derived orthotopic xenograft (PDOX) model.

**Methods:** The PDOX model was established in the left second mammary gland of a nude mouse by surgical orthotopic implantation (SOI) of the patient triple-negative MPBC PDOX models were randomized into two groups: G1, control group (n=6); G2, tumor-targeting *S. typhimurium* A1-R group (n=7, intravenous (i.v.) injection via tail vein, weekly, for two weeks). All mice were sacrificed on day 15. Tumor volume and body weight were measured one time per week.

**Results:** *S. typhimurium* A1-R arrested tumor growth compared to the control group (P = 0.016).

**Conclusion:** The results of the present study suggest that *S. typhimurium* A1-R has future clinical potential in triple-negative MPBC patients.

## INTRODUCTION

Triple-negative breast cancer (TNBC) is an aggressive and recalcitrant subtype of breast cancer characterized by lack of expression estrogen receptor (ER), progesterone receptor (PgR), and human epidermal growth factor receptor 2 (HER2). TNBC accounts for approximately 15 to 20% of all breast cancer (1, 2). The prognosis of the patients with metastatic TNBC (mTNBC) is particularly poor, due to the lack of effective targeted therapy (3).

Matrix-producing breast carcinoma (MPBC) is a rare and specialized histological type of metaplastic carcinoma (4). MPBC is usually triple-negative breast cancer and defined as an invasive breast carcinoma with a direct transition to a cartilaginous or osseous matrix with no intervening spindle-cell component (5, 6). Effective therapy for triple-negative MPBC has not been established due to its rarity (7).

Our laboratory pioneered the patient-derived orthotopic xenograft (PDOX) model by implanting patient-derived malignant tumor fragments into orthotopic sites in mice 30 years ago by establishing surgical orthotopic implantation (SOI) (8, 9). We have shown that the PDOX model is more patient-like compared to subcutaneous patient-derived xenograft (PDX) models (16). We have previously demonstrated that the PDOX model retains the histopathological/molecular and metastatic characteristics of the original tumor after SOI in mice (9-11). We previously showed eribulin regressed triple-negative MPBC PDOX mouse models (12, 13).

*Salmonella typhimurium* (*S. typhimurium*) A1-R is a facultative anaerobe. Green fluorescence protein (GFP)-labeled *S. typhimurium* A1-R (*S. typhimurium* A1-R -GFP), developed by our laboratory, has high tumor-targeting efficacy, due to its leucine–arginine auxotrophy, resulting in broad antitumor efficacy and limited adverse effects (14-16). *S. typhimurium* A1-R-GFP was able to inhibit or eradicate primary and metastatic tumors as monotherapy in nude mouse models of prostate (14, 16), breast (15, 17, 18), lung (19, 20), pancreatic (21-23), ovarian (24, 25), stomach (26), and cervical cancer (27), as well as sarcoma (28-30), melanoma (31), and glioma (32).

The development of new treatment strategies is needed for triple-negative MPBC patients. The present report demonstrates that *S. typhimurium* A1-R-GFP is an effective treatment strategy for triple-negative MPBC in a PDOX mouse model.

## MATERIAL AND METHODS

### Mouse studies

In the present study, a thymic *nu/nu* nude female mice (AntiCancer Inc, San Diego, CA), 6-7 weeks old, were used. All mice were kept in a barrier facility on a high efficacy particulate air (HEPA)-filtered rack under standard conditions of 12-h light/dark cycles. Mouse studies were performed with an AntiCancer Institutional Animal Care and Use Committee (IACUC)-protocol specially approved for the present study in accordance with the principles and procedures outlined in the National Institutes of Health Guide for the Care and Use of Animals under Assurance Number A3873-1.

### Previous establishment of the triple-negative MPBC PDOX model

A 43-year-old female patient with primary left breast cancer previously received a total mastectomy with axillary lymph node dissection at Kawasaki Medical School Hospital, Japan. The tumor was diagnosed as matrix-producing carcinoma without *BRCA* mutations. The results of immunohistostaining were as follows: ER (-), PgR (-), HER2 (-). The patient did not receive any neoadjuvant therapy. Informed consent was obtained for the patient’s tumor to be established in mouse models for studies, and PDOX studies were approved by the Institutional Ethics Committee of Kawasaki Medical School. A fresh resected tumor specimen was previously implanted subcutaneously in nude mice for establishment at AntiCancer Japan (33). The established tumors were divided into three mm^3^ fragments for orthotopic surgical implantation (SOI). A 5 mm skin incision on the left second mammary gland was made under anesthesia. The mammary gland was exposed, and a single fragment was implanted by SOI using 7-0 PDS II (polydioxanone) sutures (Ethicon, Inc., NJ, USA) (12). The wound was closed with 5-0 PDS II sutures (Ethicon, Inc., NJ, USA) (34).

### Preparation of S. typhimurium A1-R

GFP-expressing S. typhimurium A1-R (AntiCancer Inc.) were grown overnight in LB medium (Fisher Sci., Hanover Park, IL, USA) and then diluted 1:10 in LB medium. Bacteria were harvested in late-log phase, washed with PBS, and then diluted in PBS.

### Treatment protocol for the triple-negative MPBC PDOX model

The detailed experimental schema is shown in Fig. 1. The triple-negative MPBC PDOX models were randomized into two groups when the tumor volume was over 70 mm^3^: G1: untreated control; G2: *S. typhimurium* A1-R-GFP treatment (i.v., 5 × 10^7^ colony-forming units [CFU] *S. typhimurium* A1-R -GFP in 100 μl phosphate-buffered saline [PBS], weekly, two weeks). Each group comprised six and seven mice, respectively. Tumor size and body weight were measured once a week. Tumor volume was calculated using the following formula: tumor volume (mm) = length (mm) × width (mm) × width (mm) × 1/2. All mice were sacrificed on day 14.

**Figure 1.**
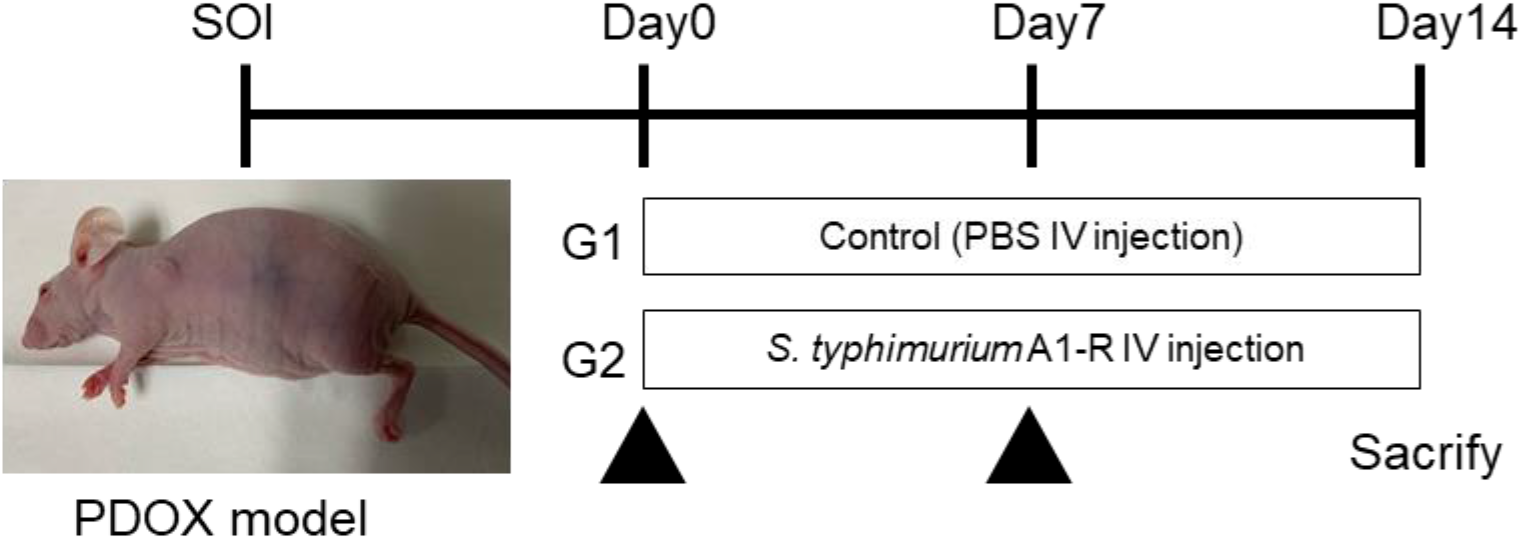
Experimental schema. *S. typhimurium* A1-R-GFP was injected into the tail vein of a MPBC TNBC PDOX nude mouse (5 × 10^7^ colony-forming units (cfu) per 100 μl of PBS). SOI: orthotopic surgical implantation. Please see Materials and Methods for details.

### Fluorescence imaging of the triple-negative MPBC PDOX

Whole-organ images were obtained and the mean fluorescence intensity of organs was determined using the UVP ChemStudio (Analytik Jena, Germany). Definition of mean fluorescence intensity: Mean raw intensity, minus mean background.

### Statistical Analysis

All statistical analyses were performed with GraphPad Prism 8.4.3 (GraphPad Software, Inc., San Diego, CA). Significant differences for comparisons of two groups were determined using the Student’s *t*-test. The experimental data are expressed as the mean ± SD. A probability value of P ≤ 0.05 was defined as statistically significant.

## RESULTS

### Selective targeting of the triple-negative MPBC PDOX by S. typhimurium A1-R -GFP

Twenty four hours after infection of triple-negative MPBC PDOX of *S. typhimurium* A1-R -GFP, the bacteria selectively targeted the tumor, as demonstrated by GFP fluorescence, as *S. typhimurium* A1-R -GFP was not detected in the liver or spleen of the triple-negative MPBC PDOX model. (Fig. 2).

**Figure 2.**
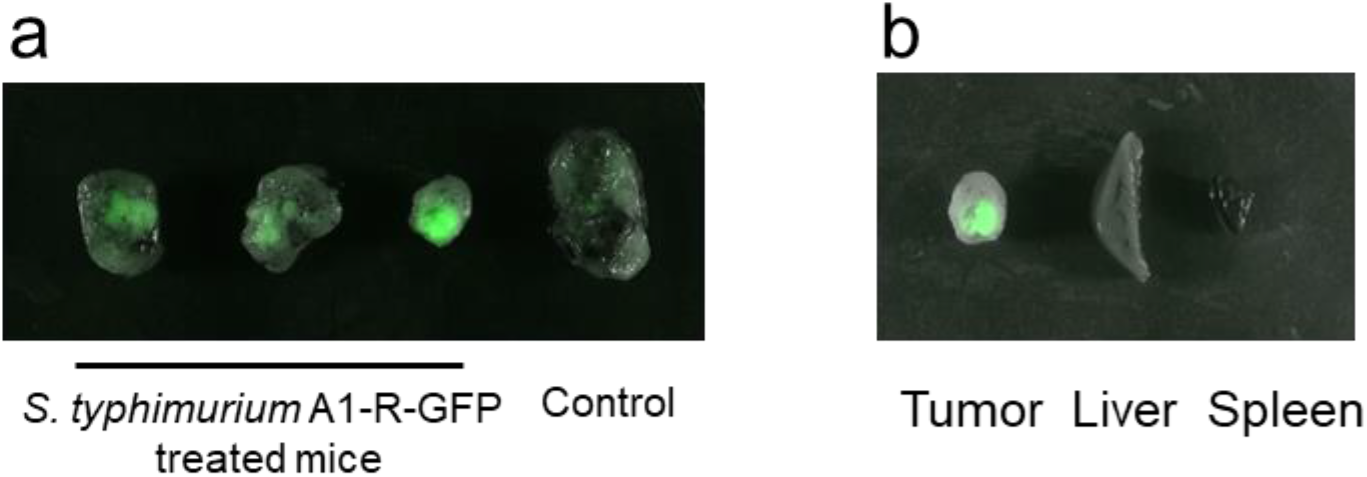
Tumor targeting by *S. typhimurium* A1-R-GFP Selective tumor targeting of S. typhimurium A1-R-GFP to the triple-negative MPBC PDOX. Iimaging was performed with the UVP ChemStudio (Analytik Jena, Germany). Please see Materials and Methods for details.

**Figure 3.**
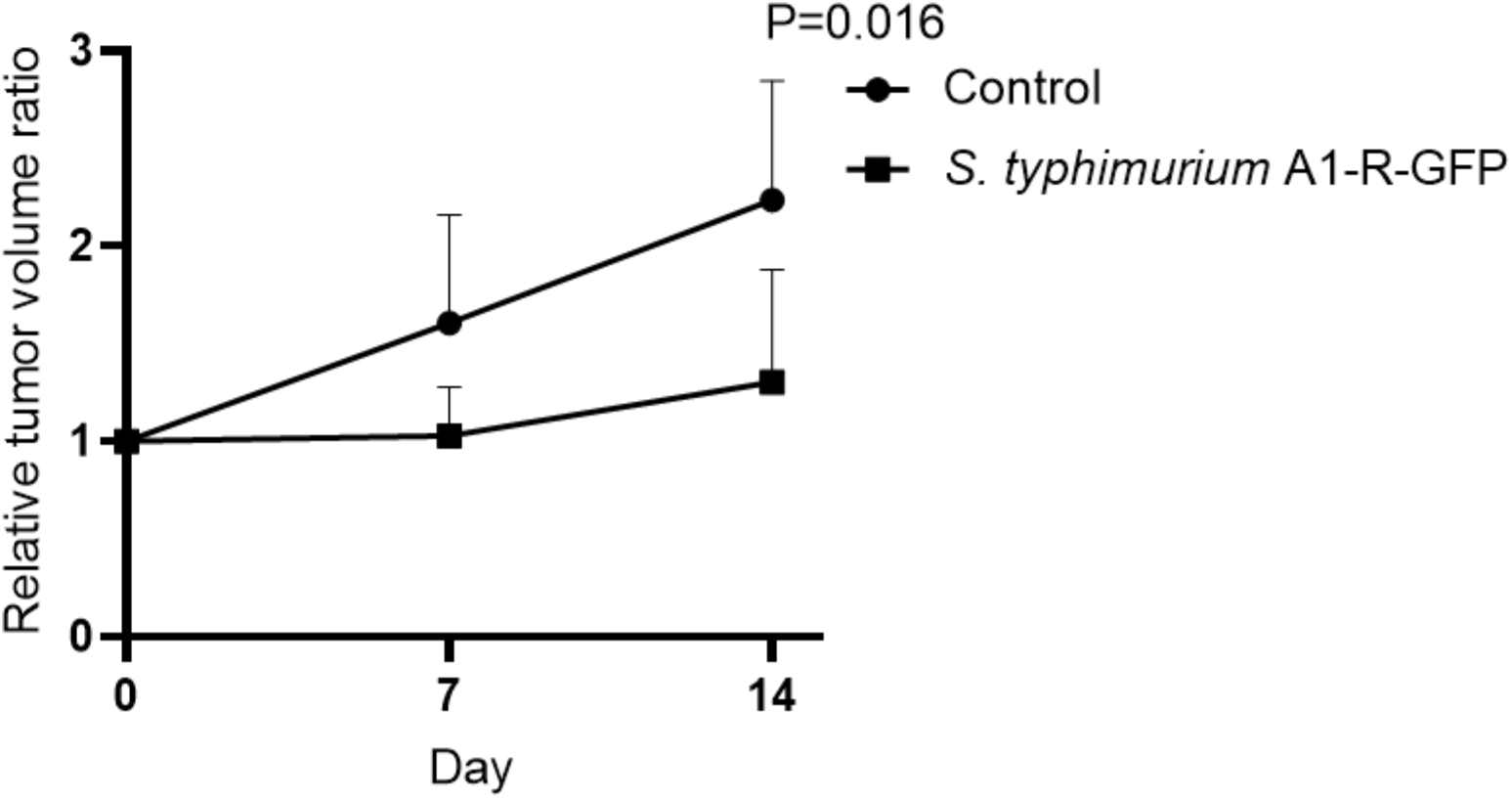
Efficacy of S. typhimurium A1-R-GFP on the triple-negative MPBC PDOX nude mice. Line graphs show the tumor volume ratio compare to day 0 from initiation of therapy. Tumors were measured at the indicated time points after the initiation of treatment. Control group n=6 vs. S. typhimurium A1-R-GFP treated group n=7. Please see materials and methods for detail. P = 0.016. Error bars represent the mean ± SD.

### S. typhimurium A1-R -GFP arrested the triple-negative MPBC PDOX

We tested the efficacy of *S. typhimurium* A1-R on the MPBC PDOX mouse model. *S. typhimurium* A1-R arrested the growth of the triple-negative MPBC (P = 0.016). The final tumor volume ratios were obtained (day 14 vs. day 0): the control group (G1) (2.236 ± 0.556); *S. typhimurium* A1-R -GFP -treated (G2) (1.303 ± 0.533).

### No effect of S. typhimurium A1-R -GFP on body weight of the triple-negative MPBC PDOX mouse model

To determine whether the drug treatments had any effect on body weight, we measured the mouse body weight at pre-treatment and post-treatment. The final body weight ratios for (day 14 vs. day 0): for the control group (G1) was 1.031 ± 0.043; and for the *S. typhimurium* A1-R -GFP -treated (G2) was 0.968 ± 0.093. There were no significant differences found in the body weight ratio between the two groups on day7, on day14 (P = 0.48, 0.73, respectively) (Fig. 4), which suggests that *S. typhimurium* A1-R -GFP had no obvious side effects.

**Figure 4.**
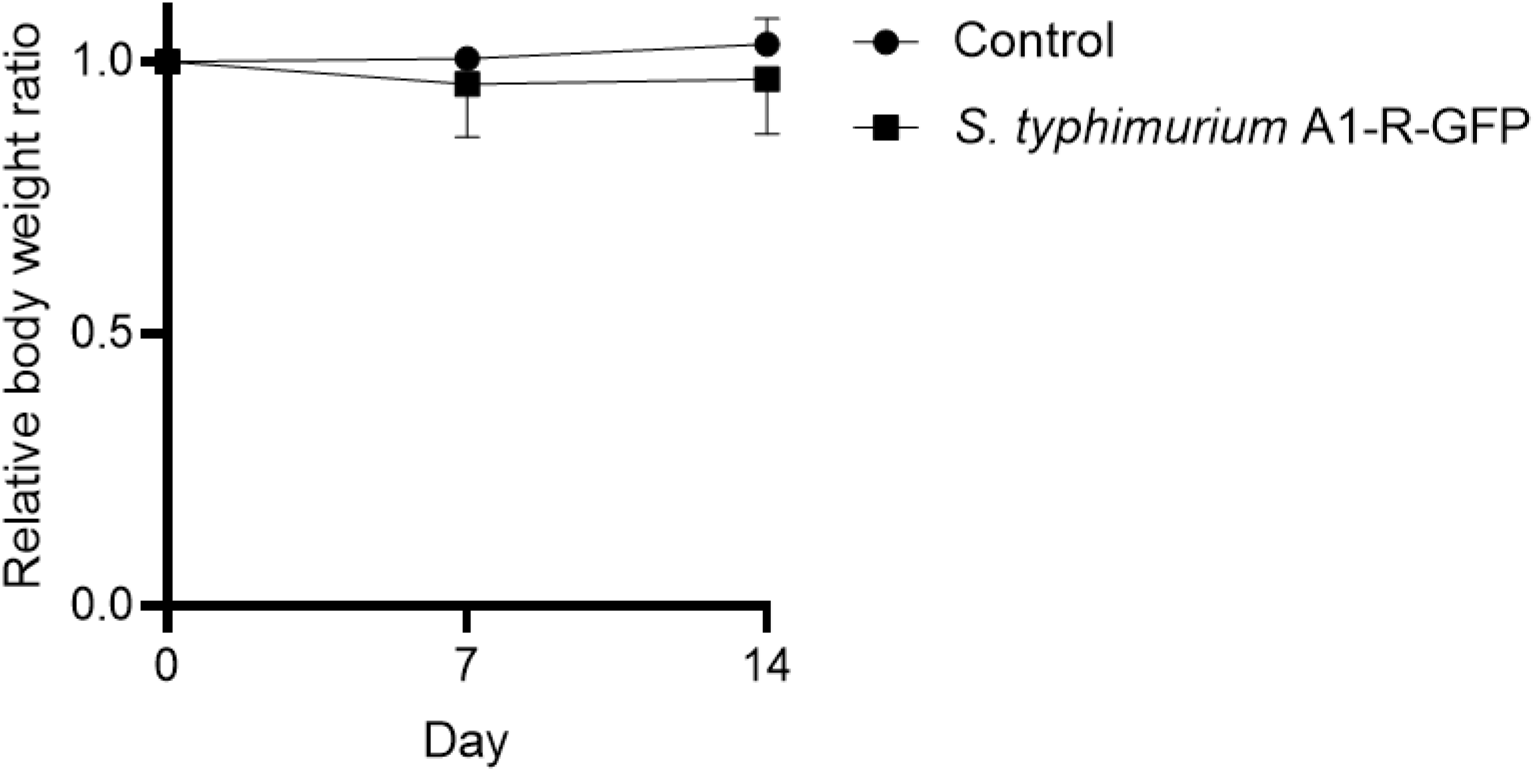
Effect of S. typhimurium A1-R-GFP on mouse body weight ratio in each group. Line graphs show the body weight ratio of mice. No significant body-weight differences were observed between the groups at the each point. Control group n=6 vs., S. typhimurium A1-R-GFP treated group n=7, on day7, on day14, P = 0.48, 0.73, respectively.

## DISCUSSION

In the present study, *S. typhimurium* A1-R-GFP selectively targeted the triple-negative MPBC PDOX and arrested its growth.

MPBC is a rare tumor with few reported studies. On only a small number of cases (35). Kusafuka et al. reported the frequency of MPBCs among all invasive breast cancer cases as only 0.2% (6). MPBC is usually TNBC and has high proliferative activity, indicated by high histological grade, high Ki-67 index, and high level of p53 expression (4, 5). Shimada et al. reported that the mean Ki-67 index of the MPBCs (45%) was higher than that of TNBCs (36%), suggesting that MPBCs are a biologically aggressive subgroup of TNBC (7).

The triple-negative MPBC is negative for ER, PR, and HER2, the type used in the present study has no approved targeted therapy. Due to its rarely, its rareness, there are only a few treatment studies for MPBC, they showed poor efficacy (7, 36, 37). Therefore, novel effective therapy is urgently needed for MPBC patients. Our previous studies of triple-negative MPBC PDOX, showed eribulin was effective (12, 13).

Bacteria therapy has gained popularity as a cancer immunotherapy in recent years (38). Salmonella, Clostridium, and other bacterial genera have been shown to control tumor growth and promote survival in animal models (38). S. typhimurium A1-R is an auxotrophic leucine-arginine facultative anaerobic S. typhimurium strain which selectively targets and proliferates in tumors of all types due to in part at least its nutritional auxotrophy which appears to be satisfied in the rich nutritional milieu of tumors, which severely restricts its growth in normal tissue (16). In the present study *S. typhimurium* A1-R-GFP selectively localized and proliferated in the triple-negative MPBC tumor. By contrast, S. *typhimurium* A1-R-GFP was undetectable in the liver, spleen. These results suggest the clinical potential of S. *typhimurium* A1-R for triple-negative MPBC.

### Conclusion

*S. typhimurium* A1-R-GFP arrested the growth of a triple-negative MPBC PDOX model without apparent toxicity. The triple-negative MPBC PDOX model should enable precise, individualized, improved therapy for patients with this recalcitrant disease.

## Acknowledgments

This paper is dedicated to the memory of A. R. Moossa, M.D., Sun Lee, M.D., Professor Li Jiaxi, and Masaki Kitajima, M.D.

## Author Contributions

K.H., J.Y., and R.M.H designed and performed experiments, analyzed data, and wrote the paper; T.M. provided tumor; C.H., Z.M., N.S., Y.S., Y.A., H.N., and M.B. gave technical support and conceptual advice. Writing, review, and/or revision of the manuscript: K.H., T.T., and R.M.H.

## REFERENCES

1 Carey LA, Perou CM, Livasy CA, Dressler LG, Cowan D, Conway K, Karaca G, Troester MA, Tse CK, Edmiston S, Deming SL, Geradts J, Cheang MCU, Nielsen TO, Moorman PG, Earp HS and Millikan RC: Race, breast cancer subtypes, and survival in the carolina breast cancer study. JAMA 295(21): 2492–2502, 2006. PMID, DOI: 10.1001/jama.295.21.2492

2 SEER Cancer Statistics Review, 1975–2014. [Internet]. National Cancer Institute. 2017.

3 Liedtke C, Mazouni C, Hess KR, Andre F, Tordai A, Mejia JA, Symmans WF, Gonzalez-Angulo AM, Hennessy B, Green M, Cristofanilli M, Hortobagyi GN and Pusztai L: Response to neoadjuvant therapy and long-term survival in patients with triple-negative breast cancer. J Clin Oncol 26(8): 1275–1281, 2008. PMID, DOI: 10.1200/jco.2007.14.4147

4 Wargotz ES and Norris HJ: Metaplastic carcinomas of the breast. I. Matrix-producing carcinoma. Hum Pathol 20(7): 628–635, 1989. PMID, DOI: 10.1016/0046-8177(89)90149-4

5 Gibson GR, Qian D, Ku JK and Lai LL: Metaplastic breast cancer: Clinical features and outcomes. Am Surg 71(9): 725–730, 2005. PMID, DOI:

6 Kusafuka K, Muramatsu K, Kasami M, Kuriki K, Hirobe K, Hayashi I, Watanabe H, Hiraki Y, Shukunami C, Mochizuki T and Kameya T: Cartilaginous features in matrix-producing carcinoma of the breast: Four cases report with histochemical and immunohistochemical analysis of matrix molecules. Mod Pathol 21(10): 1282–1292, 2008. PMID, DOI: 10.1038/modpathol.2008.120

7 Shimada K, Ishikawa T, Yamada A, Sugae S, Narui K, Shimizu D, Chishima T and Endo I: Matrix-producing carcinoma as an aggressive triple-negative breast cancer: Clinicopathological features and response to neoadjuvant chemotherapy. Anticancer Res 39(7): 3863–3869, 2019. PMID, DOI: 10.21873/anticanres.13536

8 Fu XY, Besterman JM, Monosov A and Hoffman RM: Models of human metastatic colon cancer in nude mice orthotopically constructed by using histologically intact patient specimens. Proceedings of the National Academy of Sciences 88(20): 9345–9349, 1991. PMID, DOI: 10.1073/pnas.88.20.9345

9 Hoffman RM: Patient-derived orthotopic xenografts: Better mimic of metastasis than subcutaneous xenografts. Nat Rev Cancer 15(8): 451–452, 2015. PMID, DOI: 10.1038/nrc3972

10 Furukawa T, Kubota T, Watanabe M, Kitajima M and Hoffman RM: Orthotopic transplantation of histologically intact clinical specimens of stomach cancer to nude mice: Correlation of metastatic sites in mouse and individual patient donors. Int J Cancer 53(4): 608–612, 1993. PMID, DOI: 10.1002/ijc.2910530414

11 Hoffman RM: Orthotopic metastatic mouse models for anticancer drug discovery and evaluation: A bridge to the clinic. Invest New Drugs 17(4): 343–359, 1999. PMID, DOI: 10.1023/a:1006326203858

12 Yamamoto J, Murata T, Tashiro Y, Higuchi T, Sugisawa N, Nishino H, Inubushi S, Sun YU, Lim H, Miyake K, Hongo A, Nomura T, Saitoh W, Moriya T, Tanino H, Hozumi C, Bouvet M, Singh SR, Endo I and Hoffman RM: A triple-negative matrix-producing breast carcinoma patient-derived orthotopic xenograft (pdox) mouse model is sensitive to bevacizumab and vinorelbine, regressed by eribulin and resistant to olaparib. Anticancer Res 40(5): 2509–2514, 2020. PMID, DOI: 10.21873/anticanres.14221

13 Lim HI, Yamamoto J, Inubushi S, Nishino H, Tashiro Y, Sugisawa N, Han Q, Sun YU, Choi HJ, Nam SJ, Kim MB, Lee JS, Hozumi C, Bouvet M, Singh SR and Hoffman RM: A single low dose of eribulin regressed a highly aggressive triple-negative breast cancer in a patient-derived orthotopic xenograft model. Anticancer Res 40(5): 2481–2485, 2020. PMID, DOI: 10.21873/anticanres.14218

14 Zhao M, Yang M, Li XM, Jiang P, Baranov E, Li S, Xu M, Penman S and Hoffman RM: Tumor-targeting bacterial therapy with amino acid auxotrophs of gfp-expressing salmonella typhimurium. Proc Natl Acad Sci U S A 102(3): 755–760, 2005. PMID: PMC545558, DOI: 10.1073/pnas.0408422102

15 Zhao M, Yang M, Ma H, Li X, Tan X, Li S, Yang Z and Hoffman RM: Targeted therapy with a *salmonella typhimurium* leucine-arginine auxotroph cures orthotopic human breast tumors in nude mice. Cancer Res 66(15): 7647, 2006. PMID, DOI: 10.1158/0008-5472.CAN-06-0716

16 Zhao M, Geller J, Ma H, Yang M, Penman S and Hoffman RM: Monotherapy with a tumor-targeting mutant of *salmonella typhimurium* cures orthotopic metastatic mouse models of human prostate cancer. Proceedings of the National Academy of Sciences 104(24): 10170, 2007. PMID, DOI: 10.1073/pnas.0703867104

17 Zhang Y, Tome Y, Suetsugu A, Zhang L, Zhang N, Hoffman RM and Zhao M: Determination of the optimal route of administration of salmonella typhimurium a1-r to target breast cancer in nude mice. Anticancer Res 32(7): 2501–2508, 2012. PMID, DOI:

18 Zhang Y, Miwa S, Zhang N, Hoffman RM and Zhao M: Tumor-targeting salmonella typhimurium a1-r arrests growth of breast-cancer brain metastasis. Oncotarget 6(5): 2615–2622, 2015. PMID: PMC4413605, DOI: 10.18632/oncotarget.2811

19 Uchugonova A, Zhao M, Zhang Y, Weinigel M, König K and Hoffman RM: Cancer-cell killing by engineered salmonella imaged by multiphoton tomography in live mice. Anticancer Res 32(10): 4331–4337, 2012. PMID, DOI:

20 Liu F, Zhang L, Hoffman RM and Zhao M: Vessel destruction by tumor-targeting salmonella typhimurium a1-r is enhanced by high tumor vascularity. Cell Cycle 9(22): 4518–4524, 2010. PMID: PMC3048048, DOI: 10.4161/cc.9.22.13744

21 Nagakura C, Hayashi K, Zhao M, Yamauchi K, Yamamoto N, Tsuchiya H, Tomita K, Bouvet M and Hoffman RM: Efficacy of a genetically-modified salmonella typhimurium in an orthotopic human pancreatic cancer in nude mice. Anticancer Res 29(6): 1873–1878, 2009. PMID, DOI:

22 Yam C, Zhao M, Hayashi K, Ma H, Kishimoto H, McElroy M, Bouvet M and Hoffman RM: Monotherapy with a tumor-targeting mutant of s. Typhimurium inhibits liver metastasis in a mouse model of pancreatic cancer. J Surg Res 164(2): 248–255, 2010. PMID: PMC2888721, DOI: 10.1016/j.jss.2009.02.023

23 Hiroshima Y, Zhao M, Zhang Y, Maawy A, Hassanein MK, Uehara F, Miwa S, Yano S, Momiyama M, Suetsugu A, Chishima T, Tanaka K, Bouvet M, Endo I and Hoffman RM: Comparison of efficacy of salmonella typhimurium a1-r and chemotherapy on stem-like and non-stem human pancreatic cancer cells. Cell Cycle 12(17): 2774–2780, 2013. PMID: PMC3899191, DOI: 10.4161/cc.25872

24 Matsumoto Y, Miwa S, Zhang Y, Hiroshima Y, Yano S, Uehara F, Yamamoto M, Toneri M, Bouvet M, Matsubara H, Hoffman RM and Zhao M: Efficacy of tumor-targeting salmonella typhimurium a1-r on nude mouse models of metastatic and disseminated human ovarian cancer. J Cell Biochem 115(11): 1996–2003, 2014. PMID, DOI: 10.1002/jcb.24871

25 Matsumoto Y, Miwa S, Zhang Y, Zhao M, Yano S, Uehara F, Yamamoto M, Hiroshima Y, Toneri M, Bouvet M, Matsubara H, Tsuchiya H and Hoffman RM: Intraperitoneal administration of tumor-targeting salmonella typhimurium a1-r inhibits disseminated human ovarian cancer and extends survival in nude mice. Oncotarget 6(13): 11369–11377, 2015. PMID: PMC4484462, DOI: 10.18632/oncotarget.3607

26 Yano S, Zhang Y, Zhao M, Hiroshima Y, Miwa S, Uehara F, Kishimoto H, Tazawa H, Bouvet M, Fujiwara T and Hoffman RM: Tumor-targeting salmonella typhimurium a1-r decoys quiescent cancer cells to cycle as visualized by fucci imaging and become sensitive to chemotherapy. Cell Cycle 13(24): 3958–3963, 2014. PMID: PMC4615054, DOI: 10.4161/15384101.2014.964115

27 Hiroshima Y, Zhang Y, Zhao M, Zhang N, Murakami T, Maawy A, Mii S, Uehara F, Yamamoto M, Miwa S, Yano S, Momiyama M, Mori R, Matsuyama R, Chishima T, Tanaka K, Ichikawa Y, Bouvet M, Endo I and Hoffman RM: Tumor-targeting salmonella typhimurium a1-r in combination with trastuzumab eradicates her-2-positive cervical cancer cells in patient-derived mouse models. PLoS One 10(6): e0120358, 2015. PMID: PMC4457918, DOI: 10.1371/journal.pone.0120358

28 Murakami T, DeLong J, Eilber FC, Zhao M, Zhang Y, Zhang N, Singh A, Russell T, Deng S, Reynoso J, Quan C, Hiroshima Y, Matsuyama R, Chishima T, Tanaka K, Bouvet M, Chawla S, Endo I and Hoffman RM: Tumor-targeting salmonella typhimurium a1-r in combination with doxorubicin eradicate soft tissue sarcoma in a patient-derived orthotopic xenograft (pdox) model. Oncotarget 7(11): 12783–12790, 2016. PMID: PMC4914321, DOI: 10.18632/oncotarget.7226

29 Hiroshima Y, Zhao M, Zhang Y, Zhang N, Maawy A, Murakami T, Mii S, Uehara F, Yamamoto M, Miwa S, Yano S, Momiyama M, Mori R, Matsuyama R, Chishima T, Tanaka K, Ichikawa Y, Bouvet M, Endo I and Hoffman RM: Tumor-targeting salmonella typhimurium a1-r arrests a chemo-resistant patient soft-tissue sarcoma in nude mice. PLoS One 10(8): e0134324, 2015. PMID: PMC4523197, DOI: 10.1371/journal.pone.0134324

30 Kiyuna T, Murakami T, Tome Y, Kawaguchi K, Igarashi K, Zhang Y, Zhao M, Li Y, Bouvet M, Kanaya F, Singh A, Dry S, Eilber FC and Hoffman RM: High efficacy of tumor-targeting salmonella typhimurium a1-r on a doxorubicin- and dactolisib-resistant follicular dendritic-cell sarcoma in a patient-derived orthotopic xenograft pdox nude mouse model. Oncotarget 7(22): 33046–33054, 2016. PMID: PMC5078074, DOI: 10.18632/oncotarget.8848

31 Yamamoto M, Zhao M, Hiroshima Y, Zhang Y, Shurell E, Eilber FC, Bouvet M, Noda M and Hoffman RM: Efficacy of tumor-targeting salmonella a1-r on a melanoma patient-derived orthotopic xenograft (pdox) nude-mouse model. PLoS One 11(8): e0160882, 2016. PMID: PMC4976963, DOI: 10.1371/journal.pone.0160882

32 Momiyama M, Zhao M, Kimura H, Tran B, Chishima T, Bouvet M, Endo I and Hoffman RM: Inhibition and eradication of human glioma with tumor-targeting salmonella typhimurium in an orthotopic nude-mouse model. Cell Cycle 11(3): 628–632, 2012. PMID: PMC3315098, DOI: 10.4161/cc.11.3.19116

33 Murata T, Hozumi C, Hiroshima Y, Shimoya K, Hongo A, Inubushi S, Tanino H and Hoffman RM: Co-implantation of tumor and extensive surrounding tissue improved the establishment rate of surgical specimens of human-patient cancer in nude mice: Toward the goal of universal individualized cancer therapy. In Vivo 34(6): 3241–3245, 2020. PMID, DOI: 10.21873/invivo.12160

34 Fu X, Le P and Hoffman RM: A metastatic orthotopic-transplant nude-mouse model of human patient breast cancer. Anticancer Res 13(4): 901–904, 1993. PMID, DOI:

35 Ayar S, Dyess DL and Carter E: Matrix-producing carcinoma: A rare variant of metaplastic breast carcinoma with heterologous elements. Breast J 16(4): 420–423, 2010. PMID, DOI: 10.1111/j.1524-4741.2010.00925.x

36 Al Sayed AD, El Weshi AN, Tulbah AM, Rahal MM and Ezzat AA: Metaplastic carcinoma of the breast clinical presentation, treatment results and prognostic factors. Acta Oncol 45(2): 188–195, 2006. PMID, DOI: 10.1080/02841860500513235

37 Hennessy BT, Giordano S, Broglio K, Duan Z, Trent J, Buchholz TA, Babiera G, Hortobagyi GN and Valero V: Biphasic metaplastic sarcomatoid carcinoma of the breast. Ann Oncol 17(4): 605–613, 2006. PMID, DOI: 10.1093/annonc/mdl006

38 Forbes NS, Coffin RS, Deng L, Evgin L, Fiering S, Giacalone M, Gravekamp C, Gulley JL, Gunn H, Hoffman RM, Kaur B, Liu K, Lyerly HK, Marciscano AE, Moradian E, Ruppel S, Saltzman DA, Tattersall PJ, Thorne S, Vile RG, Zhang HH, Zhou S and McFadden G: White paper on microbial anti-cancer therapy and prevention. Journal for ImmunoTherapy of Cancer 6(1): 78, 2018. PMID, DOI: 10.1186/s40425-018-0381-3

